# EXPLANe: An Extensible Framework for Poster Annotation with Mobile Devices

**DOI:** 10.1101/121178

**Authors:** Nikhil Gopal, Andrew Su, Chunlei Wu, Sean D. Mooney

## Abstract

**Summary:** Scientific posters tend to be brief, unstructured, and generally unsuitable for communication beyond a poster session. This paper describes EXPLANe, a framework for annotating posters using optical text recognition and web services on mobile devices. EXPLANe is demonstrated through an interface to the MyVariant.info variant annotation web services, and provides users a list of biological information linked with genetic variants (as found via extracted RSIDs from annotated posters). This paper delineates the architecture of the application, and includes results of a five-part evaluation we conducted. Researchers and developers can use the existing codebase as a foundation from which to generate their own annotation tabs when analyzing and annotating posters.

**Availability:** Alpha EXPLANe software is available as an open source application at https://github.com/ngopal/EXPLANe

**Contact:** Sean D. Mooney

## 1 Introduction

Although posters are a first-line medium for presentation of novel scientific research, posters constrain authors to a limited amount space. As a consequence of these space constraints, otherwise pertinent information and context are omitted from posters, requiring readers to obtain further information and context from those presenting posters.

Posters in genetics often include genetic variant annotations. Although variants are richly annotated and linked to a variety of bioinformatics resources, these annotations are primarily accessible through a desktop computer. In order to increase accessibility and timeliness of these annotation information, we provide an extensible poster annotation framework other researchers can build upon--EXPLANe. This paper demonstrates EXPLANe through a mobile application that allows end-users to take a picture of a poster containing RSID variant identifiers, submit the poster for processing, and returns linked information contained in credible resources.

Since smartphones provide a novel touch interface, are portable, and are connected to the Internet, many mobile applications taking advantage of these capabilities have been developed. Mobile applications are generally considered to be timely, contextual, and convenient [1]. Many mobile applications have been built to facilitate tasks that would otherwise require a standalone desktop, such as illustrating macromolecular structure, or analysis results of increasingly complex biological information [2]–[6]. Recently, mobile applications have also been developed to act either directly as an endpoint for query services, or using query services as part of their core functionality, shifting computation from the mobile device to a web server [7], [8]. EXPLANe is a mobile application that queries Optical Character Recognition tools (OCR) and variant annotation services to provide a summary of genetic information.

Much research has been conducted on accurately parsing gene names from scientific texts. Although there are generally accepted conventions with gene names, such as capitalizing all letters for human genes, there are a number of semantic details that complicate this task. For instance, gene name nomenclature is not standardized and genes may have multiple names [9]. In addition, gene names vary in terms of digits, letters, special characters, Greek letters, and some genes have compound names [9]. Other challenges associated with gene mapping are gene mention detection (identifying text referring to gene names), building a gene name dictionary, tokenization (providing a list of variations/synonyms/abbreviations for a gene), string matching (approximate string matching due to variability in naming), and false positive filtering (assessing whether a found gene is actually a disease, ambiguously named, etc.) [10].

A number of methods have been used to successfully extract gene names from text, with improvements and refinements of these methods over time. For example, machine learning approaches such as rule-based systems or support vector machines obtain precision and recall of about 83% and 84% respectively [13], [14]. Since the effect of features seems to be small, machine learning approaches seem to perform the best when using all available features [14]. Approaches such as GeNo and BioTagger, which use a dictionary approach in addition to machine learning, tend to yield slightly better precision and recall results of about 88% and 85%-89% respectively [10], [15]. To improve performance even further, researchers have created approaches targeting certain sub-domains (e.g. expecting genes from yeast only). LeadMine is a more recent approach, in the domain of drug name identification, uses a grammar- and dictionary-driven method has shown promising results with close to 90% precision and 85% recall, although these measures were not calculated using the BioCreAtIvE corpus [16]. Another example is pGenN, in the domain of plants, is a dictionary based approach that is able to obtain 91% precision and 87% recall [17].

Although the performance of gene name recognition methods have been steadily improving, the false positive rate remains too high to be able to use in this particular mobiles application. Thus, to minimize false positive calls to API services, and to include functionality for all organisms, EXPLANe is designed to parse and use Reference SNP cluster IDs (RSIDs) from poster text.

*Biological Web Services*: Since storage and memory capacities are limited on mobiles devices (relative to desktop computers), EXPLANe is designed to use a data federation approach, whereby required data are accessed through web services.

MyVariant.info has an application user interface (API) that is well suited for this task [18]. MyVariant.info serves as an up-to-date repository for variant information, sourcing information from more than 10 well-used bioinformatics repositories [18]. Queries may be made to MyVariant.info using RSIDs, chromosome location, and HGVS nomenclature. Additionally, MyGene.info is another endpoint allowing one to query using gene information, rather than variant information, although MyGene.info specifically is currently not queried in EXPLANe. As of July 2013, MyGene.info endpoint averaged over 3 million requests per month [18]. Furthermore, the MyVariant.info and MyGene.info endpoints are also available through an R package. However, other services, such as BioWAP, also provide access to federated biological resources and structured results, although specifically for protein-related data [19], [20].

## 2 Workflow and Development

The workflow described for the example application is implemented in Javascript/HTML5/CSS3 and transpiled to iOS and Android using Cordova/PhoneGap software. Specifically, the data binding operations are handled using D3.js [21]. Figure 1 below provides an overview of the workflow.

**Figure 1.**
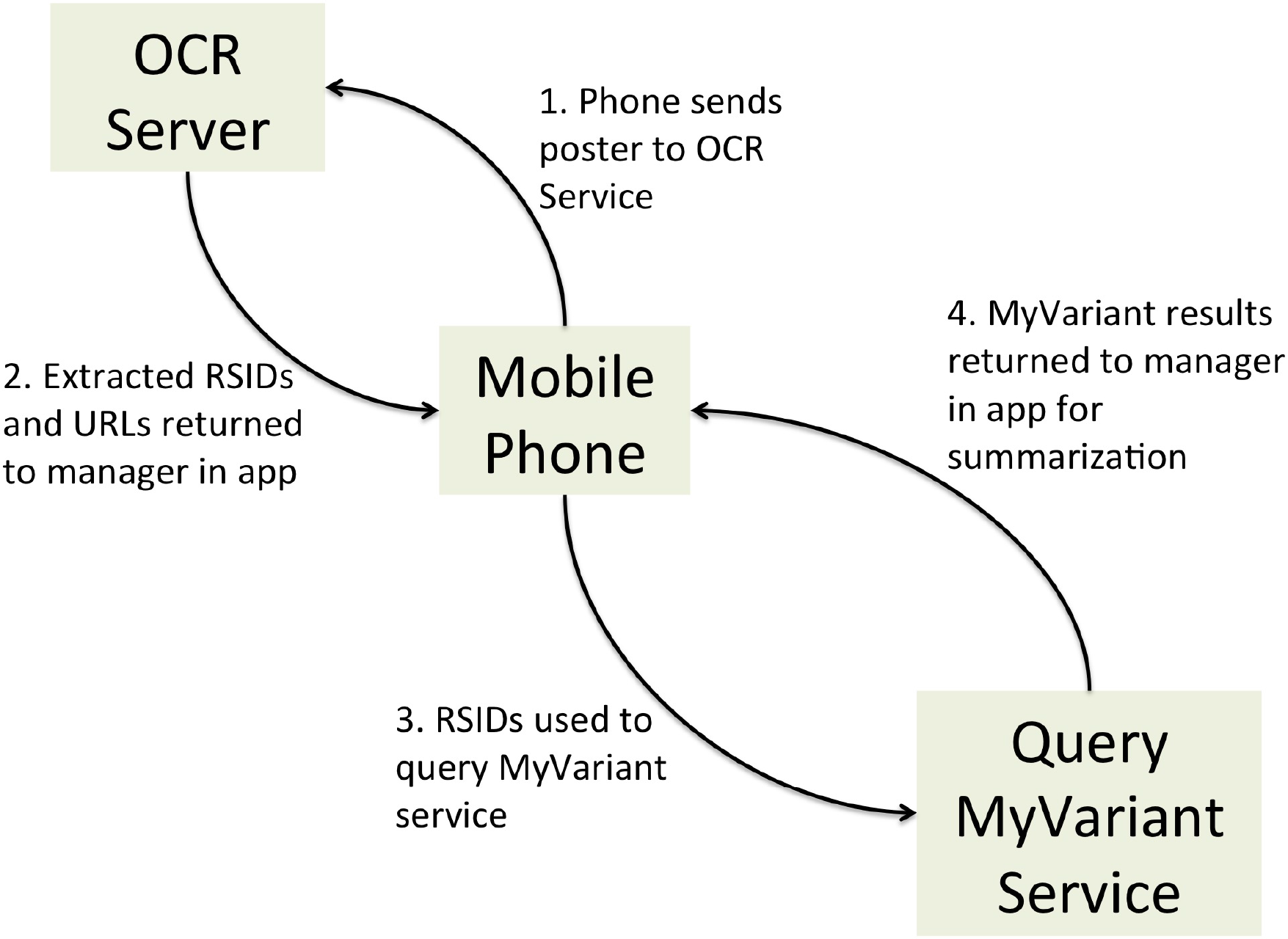
An overview of the workflow of EXPLANe. Once a user captures a picture and uploads it to the application, the application sends the poster to an OCR service that parses the text and extracts the RSIDs. Found RSIDs are returned to the application, which creates and submits a query to MyVariant.info. The MyVariant.info service returns a JSON file containing information linked to the RSID, which is summarized and displayed back to the user.

The EXPLANe framework facilitates the following workflow:

1. Mobile users provide a photograph of a poster to the application. The photograph should be of reasonably high quality so the Optical Character Recognition (OCR) service can accurately read the contents.
2. The application uploads the image to an OCR service, which returns RSIDs. Our OCR server is set to have a generous 10-minute connection timeout, which is more than sufficient to upload, process, and return RSIDs in most cases. It is possible to perform the upload step using mobile data, although we recommend connecting to Wi-Fi.
3. The RSIDs are automatically formatted into a URL and are submitted to the MyVariant.info [18] service as a query. EXPLANe is parameterized to only process up to 100 RSIDs, although this parameter may be adjust to accommodate up to 1000 RSIDs. Currently, the query is executed via a GET request for each RSID, In the future, we will switch to use POST to submit the query for multiple RSIDs in one batch.
4. The query results from MyVariant.info are returned to the application in Javascript Object Notation (JSON) format. Before further processing, for the purposes of this particular mobile application, the returned JSON file is inverted (keys in the hash become the values, and the values in the hash become the keys). Since the keys representing specific types of biological information within the returned JSON file are standardized, the JSON may be inverted. For instance, the JSON may be returned with the key ‘dbsnp’ and the key ‘clinvar’, and both of those elements may contain a value under ‘rsid’. In EXPLANe, this JSON would be inverted such that ‘rsid’ would be the key, and such that ‘dbsnp’ and ‘clinvar’ would be the values. This inversion is what allows EXPLANe to present the same data across an array of biological resources. Without inverting the JSON file, one would have to iterate through a list of resources and check to see if the entry under that resource contains the information one is interested in. By inverting the JSON file, one may call the entry directly, which would return all the relevant sources containing the desired entry.
5. The application uses the information contained in the JSON results to present a summary to the mobile users. Although the summary only presents information on RSIDs, gene names, and associated phenotypes, much more information is available for inclusion in the summary.

### OCR Service

The OCR service is provided using Tesseract and NodeJS software, and is deployed on a University of Washington production server [22], [23]. Specifically, the server software is written in Express.js, and the Tesseract software is interfaced using the “node-tesseract” javascript library. When a photograph is POSTed to the server, the server uses Tesseract via “node-tesseract.”

Although Tesseract is capable of being trained to read a variety of text, our own Tesseract server is parameterized with default settings, which means that it is optimized to recognize fairly regular blocks of text, rather than single sentences and words, or blocks of text in unexpected orientations (e.g. in a circle). The following OCR URL accepts post commands: *https://ocr.iths.org/api/photo*. The source code for a deploy-ready implementation of the OCR service is available at *https://github.com/ngopal/EXPLANe-OCR-Server*

At times, parsed RSIDs may be victim to common character misrecognitions. In order to compensate for these misrecognitions, a post-parsing filter has been implemented to catch and correct common OCR errors. This is possible thanks to the predictable structure of RSIDs (i.e. “rs” followed by a string of between 3 to 9 numeric characters). The common misrecognitions that are corrected for are: “r5” to “rs”, “l” to “1”, “z” to “2”, “o” to “0”, and “e” to “0”.

### Genetic Information

The genetic information presented to the user is obtained from MyVariant.info (http://myvariant.info). MyVariant.info is a web service that centralizes genetic information and associated metadata otherwise found across several biological resources. The returned JSON contains much more information than is currently presented to the user in the EXPLANe interface. For instance, in the scenario where gene name is to be displayed, gene name can be found across several resources. Although EXPLANe identifies variants using RSID, each variant that is found in a poster is presented using its HGVS equivalent. Currently, EXPLANe only reports RSID, HGVS ID, gene name, and associated phenotypes (as found in scientific literature).

### Application Interface

Figure 2 below presents the interface of the application. In our example, we present three tabs: “Genetics”, “URLs”, and “About”. The Genetics tab is where results from the query process are presented. The “URLs” tab presents the URLs extracted from the poster. The “About” tab is used to provide context and instructions to the user. EXPLANe allows tabs to be easily added or removed.

**Figure 2.**
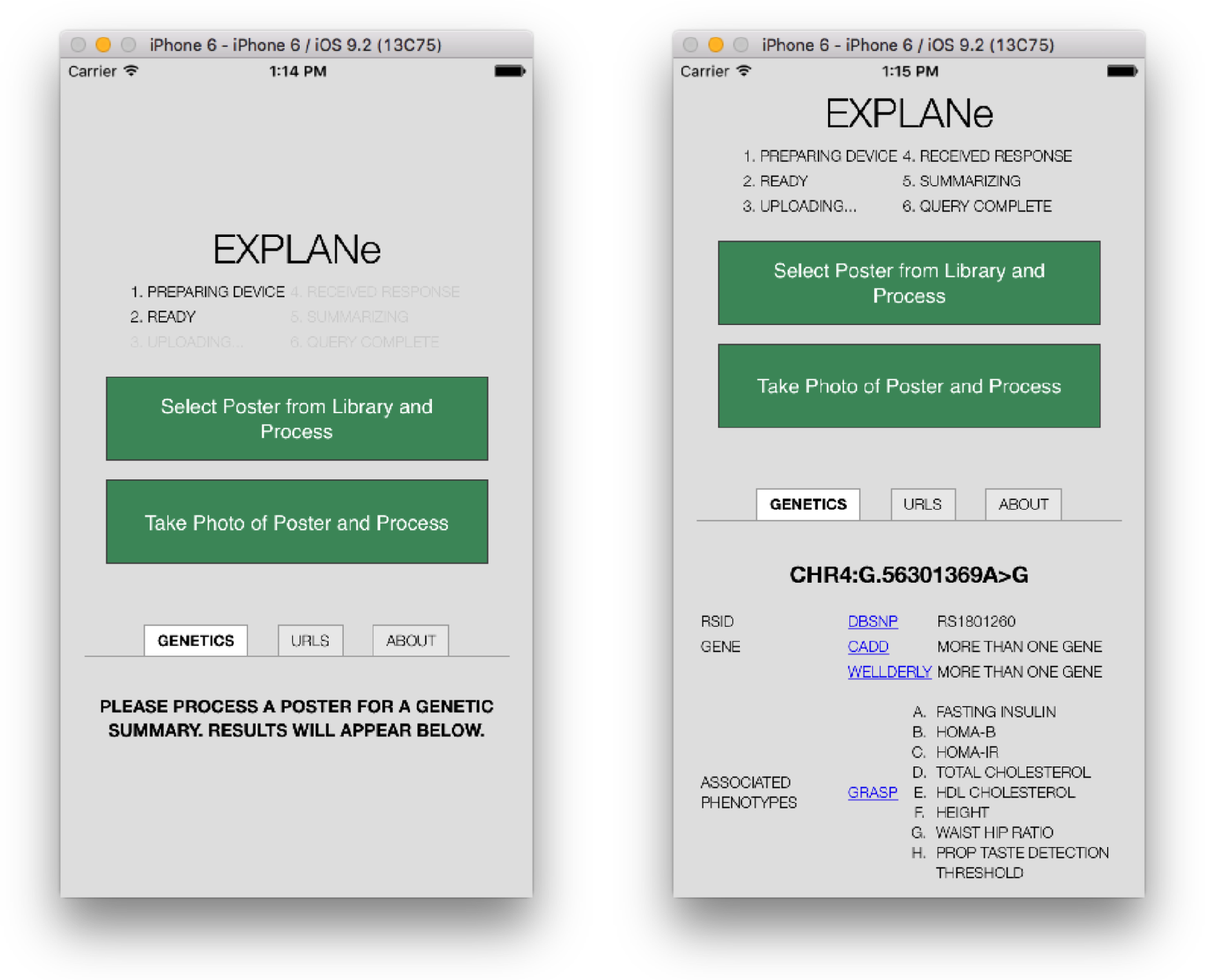
An overview of the upload interface and results interface. At the top of the application is a status bar, updating the user about the state of the application. There are six steps in the whole process. If the application is initiated successfully, the first two steps will complete and await user input (i.e. a poster image). Upon uploading, steps three through six may be completed. If the application stops at step four, then something went awry during the parsing process (and the user should be notified via an alert). Below the status bar are two large buttons the user can use to upload a picture to EXPLANe—either by uploading a pre-existing picture from the photo library on the mobile device, or taking a new poster picture on the spot (which is processed without saving the image to the photo library on the mobile device). Upon successful completion of the process, a summary is displayed under the three tabs.

## 4 Evaluation

We have evaluated EXPLANe in five different ways to understand the strengths and weaknesses of the application. We have evaluated functionality, effect of font size, effect of RSID lengths, acceptable picture distance from a poster, and consistency of repeated measures. Since the OCR server is setup with default settings, which requires blocks of text of recognition, our test posters are necessarily designed to reflect this property. Please refer to Figure 3 for an example of a test poster.

**Figure 3.**
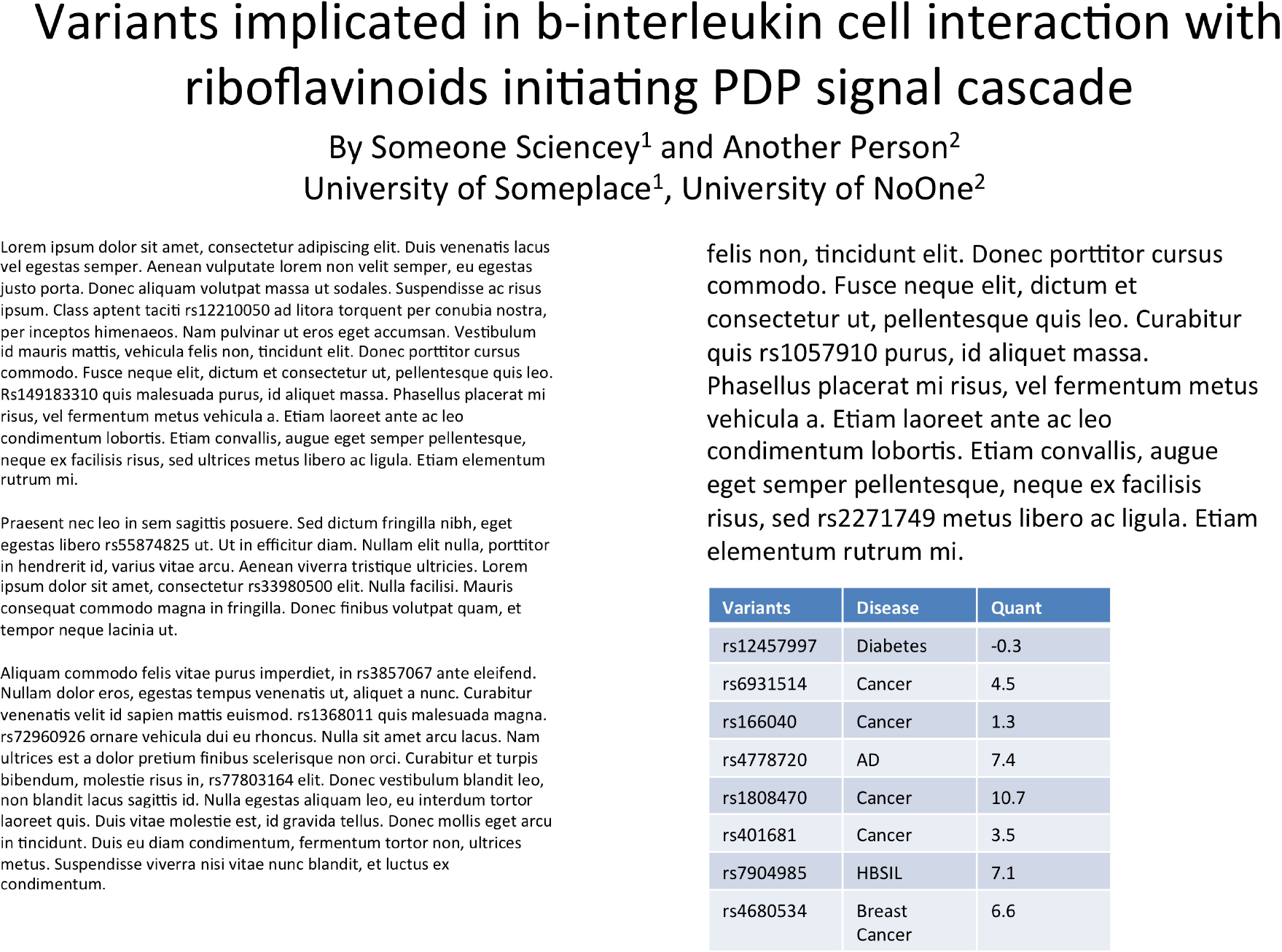
The 8.5 inch × 11inch test poster that was used for certain portions of the evaluation. This poster is representative of the type of test poster submitted to EXPLANe during the course of the evaluation.

### Evaluating Functionality

The functionality of EXPLANe was evaluated in two ways. In order to test the number of posters EXPLANe returns results for, images of various posters containing RSIDs from conferences and web resources were submitted to the mobile application. Of the 11 poster images that were submitted to EXPLANe, 3 of them (27%) returned results. Analysis of the successfully processed conference posters suggests that best results are obtained when capture angle is minimized. Of the 3 successfully processed posters, 1 of the posters contained RSIDs that were misrecognized. The ideal conditions for a poster would be minimized angles (i.e. taking a picture in as centered an orientation as possible), maximizing text size (i.e. being as close to the poster as possible), and ensuring there is sufficient lighting (for exposure), and minimizing blurs in photographs. These results are available in Table 1.

**Table 1.** Analysis results for conference posters submitted to EXPLANe. In the “Success” column, “Yes” means all RSIDs were recognized, and “No” means not all RSIDs were recognized. *Two pictures of poster 6 were available, one more distant that the other. **Poster 10 is a negative control as it did not contain any RSIDs at all.

| Poster | Success (Yes/No) | Angle |
| --- | --- | --- |
| Poster 1 | Yes | Subtle Right |
| Poster 2 | No | Subtle Top |
| Poster 3 | Yes | Subtle Left |
| Poster 4 | No | Subtly Left and Rotated |
| Poster 5 | No | Hard Top Right |
| Poster 6 | Yes | Centered |
| Poster 6 * | Yes | Centered, but distant |
| Poster 7 | Yes | Centered, but distant |
| Poster 8 | No | Subtle Top |
| Poster 9 | No | Centered |
| Poster 10 ** | No | Subtle Bottom |

### Evaluating Effect of Font Size

In order to evaluate the effect of input text font size on returned results, we created an artificial poster with text in various font sizes and submitted them to EXPLANe. The denominator in the “Number of Correctly Recognized RSIDs” column denotes the number of RSIDs in the poster, rather than the unique number of RSIDs. Pictures of these artificial posters were created in Microsoft PowerPoint, saved as large, high-resolution images, and submitted directly to EXPLANe using an iPhone simulator. Of the 3 submitted poster variations, all 3 returned results. The artificial poster variations were tested at three font sizes— results are available in Table 2. We did not test below a font size of 12, as anything smaller than that size would be too small to reasonably read.

**Table 2.** Processing results for posters of varying font size (as submitted to EXPLANe via the iPhone simulator).

| Font Size | Number of Correctly Recognized RSIDs |
| --- | --- |
| 24 | 4 / 4 (100%) |
| 18 | 5 / 5 (100%) |
| 12 | 8 / 8 (100%) |

Although RSIDs were recognized correctly 100% of the time under all tested font sizes, URLs were incorrectly recognized, and thus unable to be successfully processed and presented in the application interface. Recognizing URLs is difficult when it is conveyed in small text, as is commonly the case with citations. More challenging still is developing a regular expression pattern that encapsulates all of the required use cases and simultaneously accounts for possible modes of processing error. At the moment, EXPLANe has not implemented the complex logic required to handle URLs that are not easily recognizable and presentable, although the application can be extended to feature this capability.

### Evaluating RSID lengths

The purpose of this experiment was to determine if the length of an RSID has an effect on whether it is detected in EXPLANe. A list of human RSIDs were downloaded from UCSC Table Browser (SNP146) and organized by character length of the RSID name [24]. The character lengths of RSIDs ranged from 5-11 (including the “rs” prefix). Due to the skewed distribution of RSID name lengths (please see Table 3), we randomly selected ten RSIDs of lengths 5, 7, 9, and 11, included them in a poster, and submitted the poster to EXPLANe. Randomly selected variants were verified to return query results from MyVariant.info. Poster pictures were taken from a distance of 2 feet. The test poster in this experiment was primarily an enlarged table with nine rows of RSIDs, as well as a title that contained an RSID.

**Table 3.** An overview of the distribution of RSID name lengths

| Lengths | 5 | 6 | 7 | 8 | 9 | 10 | 11 |
| --- | --- | --- | --- | --- | --- | --- | --- |
| Frequency | 59 | 872 | 810 | 66,686 | 317,790 | 1,014,092 | 9,898,529 |

**Table 4.** RSID length versus rate of summarization (found RSIDs / included RSIDs)

| **RSID Length** | 5 | 7 | 9 | 11 |
| --- | --- | --- | --- | --- |
| **Summarization Rate** | 10/10 | 10/10 | 10/10 | 10/10 |

Based on these results, we determine that variation in RSID name length minimally affects summarization performance.

### Evaluating Distance

In this experiment, we captured pictures of posters from various distances, ranging from 1 to 10 feet away from the poster. The first poster used for this experiment was one presented at a Gordon Research Conference (GRC). The GRC poster is 48 inches × 48 inches and contains a table with RSIDs. Due to Gordon Conference rules and restrictions, we cannot provide an image of this particular test poster. The second poster was the 8.5×11 test poster in Figure 3, containing 18 RSIDs spread across text and tables.

In Table 5, it may seem peculiar that a picture taken from a distance of 1 foot yielded no results. However, since the table in the GRC poster is 4 feet × 4 feet, and since our Tesseract implementation is parameterized to recognize blocks of text, we think there may not be enough surrounding text for our OCR server to recognize RSIDs. The physical size of the RSID text in the table of the GRC poster is 0.5 inches, and the physical size of the RSID text in the table of the 8.5×11 poster is roughly 0.2 inches. If we compare the ratios of physical text size to breaking point distance, we see that GRC poster has a ratio of 0.006 and that the 8.5×11 poster has a ratio of 0.016. It may be possible to use these ratios as a heuristic to estimate optimal physical text size or optimal distance for a picture of a poster.

**Table 5.** Results for Gordon Research Conference poster

| **Distance** | 1ft | 2ft | 3ft | 4ft | 5ft | 6ft | 7ft | 8ft | 9ft | 10ft |
| --- | --- | --- | --- | --- | --- | --- | --- | --- | --- | --- |
| **Recognition** | None | Found | Found | Found | Found | Found | Found | Err | Err | None |

**Table 6.** Results for 8.5 × 11 inch poster

| **Distance** | 1ft | 2ft | 3ft |
| --- | --- | --- | --- |
| **Recognition** | 18 / 18 | None | None |

### Evaluating Repeated Measures

In this experiment, we took several pictures of the same poster from the same distance (2 feet away). Our goal for this experiment is to determine how often results are repeated. We took five pictures of the same 8.5×11 poster where the poster was fully in frame. There were a total of 18 RSIDs contained in the poster, although only 10 are examined here.

As Table 7 shows, 12 of 18 RSIDs were recognized every time, with the remaining 6 being recognized between 3 and 4 times of the 5. No RSIDs in the poster were misrecognized. After assessing the location of the RSIDs that occurred at a frequency of 3 and 4, we could not determine any systematic source to account for the imperfect performance.

**Table 7.** Results for repeated measure of the same poster

| **RSID** | **Frequency of summarization** |
| --- | --- |
| Rs77803164 | 5 |
| Rs72960926 | 5 |
| Rs6931514 | 5 |
| Rs4778720 | 5 |
| Rs4680534 | 5 |
| Rs401681 | 5 |
| Rs3857067 | 5 |
| Rs2271749 | 5 |
| Rs166040 | 5 |
| Rs149183310 | 5 |
| Rs1368011 | 5 |
| Rs12457997 | 5 |
| Rs7904985 | 4 |
| Rs33980500 | 4 |
| Rs1808470 | 4 |
| Rs1057910 | 4 |
| Rs55874825 | 3 |
| Rs12210050 | 3 |

## 4 Extensibility

EXPLANe can be extended in a number of ways. As seen in Figure 1, the interface organizes the annotation results by categories (e.g. variant information, URLs, etc.). Adding or customizing tabs is the easiest way to use and extend the framework (full implementation instructions accompanied with code). Future work may include extending EXPLANe to include the BioWAP resource, as this would connect the OCR capability with additional protein-related resources that may provide a richer user experience. Furthermore, if the error rate of named entity recognitions is deemed tolerable, EXPLANe may be extended to recognize genes and query MyGene.info, providing a wider range of recognizable words with attached information. We anticipate that named entity recognition for gene names from scientific texts will improve over time, and that EXPLANe may eventually be updated to also process gene names. In addition to improvements in NER, performance of EXPLANe may conceivably be improved through training the Tesseract software on a scientific corpus (akin to the type of posters the OCR system will be processing), and through providing Tesseract a data dictionary containing domain-specific vocabulary terms that have been historically difficult to process, such as human gene names.

The application works very well under simulated conditions, but begins to have imperfect performance when pictures are taken of physical posters. From this observation, we infer that performance may vary depending on the characteristics and features of the mobile device being used. In certain cases, this may mean taking a picture in “landscape” mode rather than “portrait”. An ideal poster image would minimized angles (i.e. taking a picture in as centered an orientation as possible), maximize text size (i.e. being as close to the poster as possible), ensure adequate lighting (for exposure), and minimize blurs.

Overall, the capability of EXPLANe is bounded by OCR recognition performance, and the content contained in MyVariant.info. However, as mentioned above, the OCR service we have deployed is not optimized, and MyVariant.info is ever-growing in terms of supported biological resources.

## 5 Conclusion

EXPLANe provides two contributions. First, it provides an extensible framework for poster annotation that any researcher can extend, or build upon. Second, it enables readers of posters to enrich their experience with supplemental poster information in real-time. Future work may entail adding social media capability, data downloads, emailing results, and user annotation. EXPLANe is readily extensible by anyone interested in downloading the source code and standing up their own instance. Performance may be improved by optimizing Tesseract to recognize RSIDs.

